# The *Rhinolophus affinis* bat ACE2 and multiple animal orthologs are functional receptors for bat coronavirus RaTG13 and SARS-CoV-2

**DOI:** 10.1101/2020.11.16.385849

**Authors:** Pei Li, Ruixuan Guo, Yan Liu, Yintgtao Zhang, Jiaxin Hu, Xiuyuan Ou, Dan Mi, Ting Chen, Zhixia Mu, Yelin Han, Zhewei Cui, Leiliang Zhang, Xinquan Wang, Zhiqiang Wu, Jianwei Wang, Qi Jin, Zhaohui Qian

## Abstract

Bat coronavirus (CoV) RaTG13 shares the highest genome sequence identity with SARS-CoV-2 among all known coronaviruses, and also uses human angiotensin converting enzyme 2 (hACE2) for virus entry. Thus, SARS-CoV-2 is thought to have originated from bat. However, whether SARS-CoV-2 emerged from bats directly or through an intermediate host remains elusive. Here, we found that *Rhinolophus affinis* bat ACE2 (RaACE2) is an entry receptor for both SARS-CoV-2 and RaTG13, although RaACE2 binding to the receptor binding domain (RBD) of SARS-CoV-2 is markedly weaker than that of hACE2. We further evaluated the receptor activities of ACE2s from additional 16 diverse animal species for RaTG13, SARS-CoV, and SARS-CoV-2 in terms of S protein binding, membrane fusion, and pseudovirus entry. We found that the RaTG13 spike (S) protein is significantly less fusogenic than SARS-CoV and SARS-CoV-2, and seven out of sixteen different ACE2s function as entry receptors for all three viruses, indicating that all three viruses might have broad host rages. Of note, RaTG13 S pseudovirions can use mouse, but not pangolin ACE2, for virus entry, whereas SARS-CoV-2 S pseudovirions can use pangolin, but limited for mouse, ACE2s enter cells. Mutagenesis analysis revealed that residues 484 and 498 in RaTG13 and SARS-CoV-2 S proteins play critical roles in recognition of mouse and human ACE2. Finally, two polymorphous *Rhinolophous sinicus* bat ACE2s showed different susceptibilities to virus entry by RaTG13 and SARS-CoV-2 S pseudovirions, suggesting possible coevolution. Our results offer better understanding of the mechanism of coronavirus entry, host range, and virus-host coevolution.

## Introduction

Coronavirus disease 2019 (COVID-19) is caused by a newly emerged coronavirus, severe acute respiratory syndrome coronavirus 2 (SARS-CoV-2), first identified in late 2019 in Wuhan, China^1-4^, and currently it has spread to over 200 countries. On March 11, the World Health Organization declared a global pandemic of COVID-19. As of August 23rd, there are more than 23 million confirmed cases and over 800,000 deaths caused by SARS-CoV-2 worldwide^5^.

Phylogenetically, coronaviruses (CoVs) are classified into four genera, alpha, beta, gamma, and delta, and beta-CoVs are further divided into four lineages, A, B, C, and D. SARS-CoV-2 is a lineage B beta-CoV, including SARS-CoV and bat SARS-like CoVs (SL-CoV)^6,7^. The genome of SARS-CoV-2 shares approximately 80% and 96.2% nucleotide sequence identity with SARS-CoV and bat SL-CoV RaTG13, respectively^3^. The high sequence homology between SARS-CoV-2 and bat SL-CoVs suggests that SARS-CoV-2 might originate from bats^3,8,9^. However, whether zoonotic transmission from bats to humans is direct or through an intermediate animal host remains to be determined.

CoVs use their trimeric spike (S) glycoproteins to bind the receptors and mediate virus entry, and the interaction between the S protein and its cognate receptor largely determines the virus host range and tissue tropism. The S protein contains two subunits, S1 and S2. While S1 binds to the receptor, S2 contains the membrane fusion machinery. Recently we and others showed that SARS-CoV-2 uses human angiotensin converting enzyme 2 (hACE2) as the entry receptor^3,10,11^. The structure of hACE2 and the SARS-CoV-2 S protein or receptor binding domain (RBD) complex was also solved recently^12-15^, and there are extensive interactions between the SARS-CoV-2 S protein and hACE2, including 17 residues in the S protein and 20 residues in hACE2 (Table 1). Several critical residues, such as K31 and K353 in hACE2 and F486 and Q498 in the S protein, were also identified. Many animals, including cats, ferrets, minks, tigers, hamsters, dogs in lesser degree, are susceptible to SARS-CoV-2 infection^16-22^, indicating the potential broad host range of SARS-CoV-2.

**Table 1.** Alignment of critical S protein-interacting residues of different animal ACE2 proteins.

| Species<sup>o</sup> | 24<sup>o</sup> | 27<sup>o</sup> | 28<sup>o</sup> | 30<sup>o</sup> | 31<sup>o</sup> | 34<sup>o</sup> | 35<sup>o</sup> | 37<sup>o</sup> | 38<sup>o</sup> | 41<sup>o</sup> | 42<sup>o</sup> | 79<sup>o</sup> | 82<sup>o</sup> | 83<sup>o</sup> | 330<sup>o</sup> | 353<sup>o</sup> | 354<sup>o</sup> | 355<sup>o</sup> | 357<sup>o</sup> | 393<sup>o</sup> | Accession<sup>o</sup> |
| --- | --- | --- | --- | --- | --- | --- | --- | --- | --- | --- | --- | --- | --- | --- | --- | --- | --- | --- | --- | --- | --- |
| Human<sup>o</sup> | Q<sup>o</sup> | T<sup>o</sup> | F<sup>o</sup> | D<sup>o</sup> | K<sup>o</sup> | H<sup>o</sup> | E<sup>o</sup> | E<sup>o</sup> | D<sup>o</sup> | Y<sup>o</sup> | Q<sup>o</sup> | L<sup>o</sup> | M<sup>o</sup> | Y<sup>o</sup> | N<sup>o</sup> | K<sup>o</sup> | G<sup>o</sup> | D<sup>o</sup> | R<sup>o</sup> | R<sup>o</sup> | NP_068576<sup>o</sup> |
| Squirrel<sup>o</sup> | L<sup>o</sup> | T<sup>o</sup> | F<sup>o</sup> | D<sup>o</sup> | K<sup>o</sup> | Q<sup>o</sup> | E<sup>o</sup> | E<sup>o</sup> | D<sup>o</sup> | H<sup>o</sup> | Q<sup>o</sup> | L<sup>o</sup> | D<sup>o</sup> | Y<sup>o</sup> | N<sup>o</sup> | K<sup>o</sup> | G<sup>o</sup> | D<sup>o</sup> | R<sup>o</sup> | R<sup>o</sup> | XP_026252505<sup>o</sup> |
| Pangolin<sup>o</sup> | E<sup>o</sup> | T<sup>o</sup> | F<sup>o</sup> | E<sup>o</sup> | K<sup>o</sup> | S<sup>o</sup> | E<sup>o</sup> | E<sup>o</sup> | E<sup>o</sup> | Y<sup>o</sup> | Q<sup>o</sup> | I<sup>o</sup> | N<sup>o</sup> | Y<sup>o</sup> | N<sup>o</sup> | K<sup>o</sup> | H<sup>o</sup> | D<sup>o</sup> | R<sup>o</sup> | R<sup>o</sup> | XP_017505746<sup>o</sup> |
| Fox<sup>o</sup> | L<sup>o</sup> | T<sup>o</sup> | F<sup>o</sup> | E<sup>o</sup> | K<sup>o</sup> | Y<sup>o</sup> | E<sup>o</sup> | E<sup>o</sup> | E<sup>o</sup> | Y<sup>o</sup> | Q<sup>o</sup> | L<sup>o</sup> | T<sup>o</sup> | Y<sup>o</sup> | N<sup>o</sup> | K<sup>o</sup> | G<sup>o</sup> | D<sup>o</sup> | R<sup>o</sup> | R<sup>o</sup> | XP_025842512<sup>o</sup> |
| Civet<sup>o</sup> | L<sup>o</sup> | T<sup>o</sup> | F<sup>o</sup> | E<sup>o</sup> | T<sup>o</sup> | Y<sup>o</sup> | E<sup>o</sup> | Q<sup>o</sup> | E<sup>o</sup> | Y<sup>o</sup> | Q<sup>o</sup> | L<sup>o</sup> | T<sup>o</sup> | Y<sup>o</sup> | N<sup>o</sup> | K<sup>o</sup> | G<sup>o</sup> | D<sup>o</sup> | R<sup>o</sup> | R<sup>o</sup> | AAX63775<sup>o</sup> |
| Camel<sup>o</sup> | L<sup>o</sup> | T<sup>o</sup> | F<sup>o</sup> | E<sup>o</sup> | E<sup>o</sup> | H<sup>o</sup> | E<sup>o</sup> | E<sup>o</sup> | D<sup>o</sup> | Y<sup>o</sup> | Q<sup>o</sup> | T<sup>o</sup> | T<sup>o</sup> | Y<sup>o</sup> | N<sup>o</sup> | K<sup>o</sup> | G<sup>o</sup> | D<sup>o</sup> | R<sup>o</sup> | R<sup>o</sup> | XP_006194263<sup>o</sup> |
| Ferret<sup>o</sup> | L<sup>o</sup> | T<sup>o</sup> | F<sup>o</sup> | E<sup>o</sup> | K<sup>o</sup> | Y<sup>o</sup> | E<sup>o</sup> | E<sup>o</sup> | E<sup>o</sup> | Y<sup>o</sup> | Q<sup>o</sup> | H<sup>o</sup> | T<sup>o</sup> | Y<sup>o</sup> | N<sup>o</sup> | K<sup>o</sup> | R<sup>o</sup> | D<sup>o</sup> | R<sup>o</sup> | R<sup>o</sup> | NP_001297119<sup>o</sup> |
| Rat<sup>o</sup> | K<sup>o</sup> | S<sup>o</sup> | F<sup>o</sup> | N<sup>o</sup> | K<sup>o</sup> | Q<sup>o</sup> | E<sup>o</sup> | E<sup>o</sup> | D<sup>o</sup> | Y<sup>o</sup> | Q<sup>o</sup> | I<sup>o</sup> | N<sup>o</sup> | F<sup>o</sup> | N<sup>o</sup> | H<sup>o</sup> | G<sup>o</sup> | D<sup>o</sup> | R<sup>o</sup> | R<sup>o</sup> | NP_001012006<sup>o</sup> |
| Mouse<sup>o</sup> | N<sup>o</sup> | T<sup>o</sup> | F<sup>o</sup> | N<sup>o</sup> | N<sup>o</sup> | Q<sup>o</sup> | E<sup>o</sup> | E<sup>o</sup> | D<sup>o</sup> | Y<sup>o</sup> | Q<sup>o</sup> | T<sup>o</sup> | S<sup>o</sup> | F<sup>o</sup> | N<sup>o</sup> | H<sup>o</sup> | G<sup>o</sup> | D<sup>o</sup> | R<sup>o</sup> | R<sup>o</sup> | NP_001123985<sup>o</sup> |
| Pig<sup>o</sup> | L<sup>o</sup> | T<sup>o</sup> | F<sup>o</sup> | E<sup>o</sup> | K<sup>o</sup> | L<sup>o</sup> | E<sup>o</sup> | E<sup>o</sup> | D<sup>o</sup> | Y<sup>o</sup> | Q<sup>o</sup> | I<sup>o</sup> | T<sup>o</sup> | Y<sup>o</sup> | N<sup>o</sup> | K<sup>o</sup> | G<sup>o</sup> | D<sup>o</sup> | R<sup>o</sup> | R<sup>o</sup> | NP_001116542<sup>o</sup> |
| RA Bat<sup>o</sup> | R<sup>o</sup> | I<sup>o</sup> | F<sup>o</sup> | D<sup>o</sup> | N<sup>o</sup> | H<sup>o</sup> | E<sup>o</sup> | E<sup>o</sup> | D<sup>o</sup> | Y<sup>o</sup> | Q<sup>o</sup> | L<sup>o</sup> | N<sup>o</sup> | Y<sup>o</sup> | N<sup>o</sup> | K<sup>o</sup> | G<sup>o</sup> | D<sup>o</sup> | R<sup>o</sup> | R<sup>o</sup> | QMQ39244<sup>o</sup> |
| RS-YN Bat<sup>o</sup> | E<sup>o</sup> | M<sup>o</sup> | F<sup>o</sup> | D<sup>o</sup> | K<sup>o</sup> | T<sup>o</sup> | K<sup>o</sup> | E<sup>o</sup> | D<sup>o</sup> | H<sup>o</sup> | Q<sup>o</sup> | L<sup>o</sup> | N<sup>o</sup> | Y<sup>o</sup> | N<sup>o</sup> | K<sup>o</sup> | G<sup>o</sup> | D<sup>o</sup> | R<sup>o</sup> | R<sup>o</sup> | AGZ48803<sup>o</sup> |
| RS-HB Bat<sup>o</sup> | R<sup>o</sup> | T<sup>o</sup> | F<sup>o</sup> | D<sup>o</sup> | E<sup>o</sup> | S<sup>o</sup> | E<sup>o</sup> | E<sup>o</sup> | N<sup>o</sup> | Y<sup>o</sup> | Q<sup>o</sup> | L<sup>o</sup> | N<sup>o</sup> | Y<sup>o</sup> | N<sup>o</sup> | K<sup>o</sup> | G<sup>o</sup> | D<sup>o</sup> | R<sup>o</sup> | R<sup>o</sup> | ADN93475<sup>o</sup> |
| Guinea pig<sup>o</sup> | L<sup>o</sup> | I<sup>o</sup> | F<sup>o</sup> | D<sup>o</sup> | E<sup>o</sup> | S<sup>o</sup> | E<sup>o</sup> | E<sup>o</sup> | N<sup>o</sup> | Y<sup>o</sup> | Q<sup>o</sup> | L<sup>o</sup> | N<sup>o</sup> | Y<sup>o</sup> | N<sup>o</sup> | K<sup>o</sup> | N<sup>o</sup> | D<sup>o</sup> | R<sup>o</sup> | R<sup>o</sup> | ACT66270<sup>o</sup> |
| Deer<sup>o</sup> | Q<sup>o</sup> | T<sup>o</sup> | F<sup>o</sup> | E<sup>o</sup> | K<sup>o</sup> | H<sup>o</sup> | E<sup>o</sup> | E<sup>o</sup> | D<sup>o</sup> | Y<sup>o</sup> | Q<sup>o</sup> | M<sup>o</sup> | T<sup>o</sup> | Y<sup>o</sup> | N<sup>o</sup> | K<sup>o</sup> | G<sup>o</sup> | D<sup>o</sup> | R<sup>o</sup> | R<sup>o</sup> | XP_020768965<sup>o</sup> |
| Hedgehog<sup>o</sup> | Q<sup>o</sup> | S<sup>o</sup> | F<sup>o</sup> | T<sup>o</sup> | T<sup>o</sup> | N<sup>o</sup> | E<sup>o</sup> | E<sup>o</sup> | N<sup>o</sup> | Y<sup>o</sup> | Q<sup>o</sup> | L<sup>o</sup> | K<sup>o</sup> | F<sup>o</sup> | K<sup>o</sup> | L<sup>o</sup> | N<sup>o</sup> | D<sup>o</sup> | R<sup>o</sup> | R<sup>o</sup> | XP_004710002<sup>o</sup> |
| Koala<sup>o</sup> | R<sup>o</sup> | E<sup>o</sup> | F<sup>o</sup> | E<sup>o</sup> | T<sup>o</sup> | K<sup>o</sup> | E<sup>o</sup> | E<sup>o</sup> | E<sup>o</sup> | Y<sup>o</sup> | Q<sup>o</sup> | I<sup>o</sup> | T<sup>o</sup> | F<sup>o</sup> | N<sup>o</sup> | K<sup>o</sup> | G<sup>o</sup> | D<sup>o</sup> | R<sup>o</sup> | R<sup>o</sup> | XP_020863153<sup>o</sup> |
| Turtle<sup>o</sup> | E<sup>o</sup> | N<sup>o</sup> | F<sup>o</sup> | S<sup>o</sup> | E<sup>o</sup> | V<sup>o</sup> | Q<sup>o</sup> | E<sup>o</sup> | D<sup>o</sup> | Y<sup>o</sup> | A<sup>o</sup> | N<sup>o</sup> | K<sup>o</sup> | Y<sup>o</sup> | N<sup>o</sup> | K<sup>o</sup> | K<sup>o</sup> | D<sup>o</sup> | R<sup>o</sup> | R<sup>o</sup> | XP_006122891<sup>o</sup> |
| Snake<sup>o</sup> | V<sup>o</sup> | K<sup>o</sup> | F<sup>o</sup> | E<sup>o</sup> | Q<sup>o</sup> | A<sup>o</sup> | R<sup>o</sup> | T<sup>o</sup> | D<sup>o</sup> | Y<sup>o</sup> | N<sup>o</sup> | N<sup>o</sup> | M<sup>o</sup> | F<sup>o</sup> | N<sup>o</sup> | K<sup>o</sup> | E<sup>o</sup> | D<sup>o</sup> | R<sup>o</sup> | R<sup>o</sup> | ETE61880<sup>o</sup> |

Green: residues homologous to that of human ACE2
Orange: residues different from that of human ACE2
Black: residues identical to that of human ACE2

RaTG13 was first discovered in the *Rhinolophus affinis* bat^3^, and it can use hACE2 for virus entry^13,23^. CryoEM structure of its S protein in prefusion conformation was also solved, and all three monomers in trimeric S proteins are in “down” position^24^, revealing more stable in native conformation and significantly lower affinity to hACE2 than SARS-CoV-2 S protein. Recently, Li *et al* reported that SARS-CoV-2 and RaTG13 can use several domesticated animal orthologs of hACE2 for virus entry^23^. However, whether RaACE2 is a functional receptor for RaTG13 and SARS-CoV-2 remains unknown. In this study, we determined the susceptibility of 17 diverse animal species including *Rhinolophus affinis* to SARS-CoV-2 and RaTG13 viruses by using their S pseudovirions, and found that RaACE2 and several other ACE2s could efficiently mediate the entry of SARS-CoV-2, SARS-CoV, and RaTG13 virus. We further identified two residues, 484 and 498, that are critical for recognition of mouse and human ACE2s

## Results

To investigate the potential intermediate host for SARS-CoV-2, we determined the receptor usage and host range of RaTG13 using a pseudotype system. We also included the S protein of ZC45 in our study, sharing approximately 88% of genome nucleotide sequence identity with that of SARS-CoV-2 ^2,25^. Previously we found that removal of a conserved ER-retention motif, KxHxx, increased the level of S protein present on cell surface and incorporation into lentiviral pseudovirions^10,26^. Sequence alignment of the S proteins of SARS-CoV, SARS-CoV-2, RaTG13, and ZC45 revealed that KxHxx motif was also present on the S proteins of RaTG13 and ZC45 (Fig 1A).The last 19 amino acids of the S proteins of RaTG13 and ZC45 were removed and a 3xFLAG tag was also added to C-terminus of S proteins for detection. The plasmids encoding the S proteins of RaTG13 and ZC45 were transfected into 293T cells, and the levels of S protein expression were evaluated by western blot using various antibodies. The S proteins of RaTG13 and ZC45 were expressed at levels similar to those of SARS-CoV and SARS-CoV-2, and they were readily detected by monoclonal anti-FLAG M2 antibody (Fig 1B) and polyclonal anti-SARS-CoV-2 S2 antibodies (Supplementary Fig 1A), suggesting that the immunoepitope(s) for anti-SARS-CoV-2 S2 antibodies were also conserved among all four CoVs. The S proteins of SARS-CoV-2 and RaTG13 were also detected by anti-SARS-CoV-2 RBD antibodies, but weakly bound to rabbit polyclonal anti-SARS S1 antibody T62 and mouse monoclonal anti-SARS S1 antibody MM02 (Supplementary Figs 1B, 1C, and 1D).

**Figure 1.**
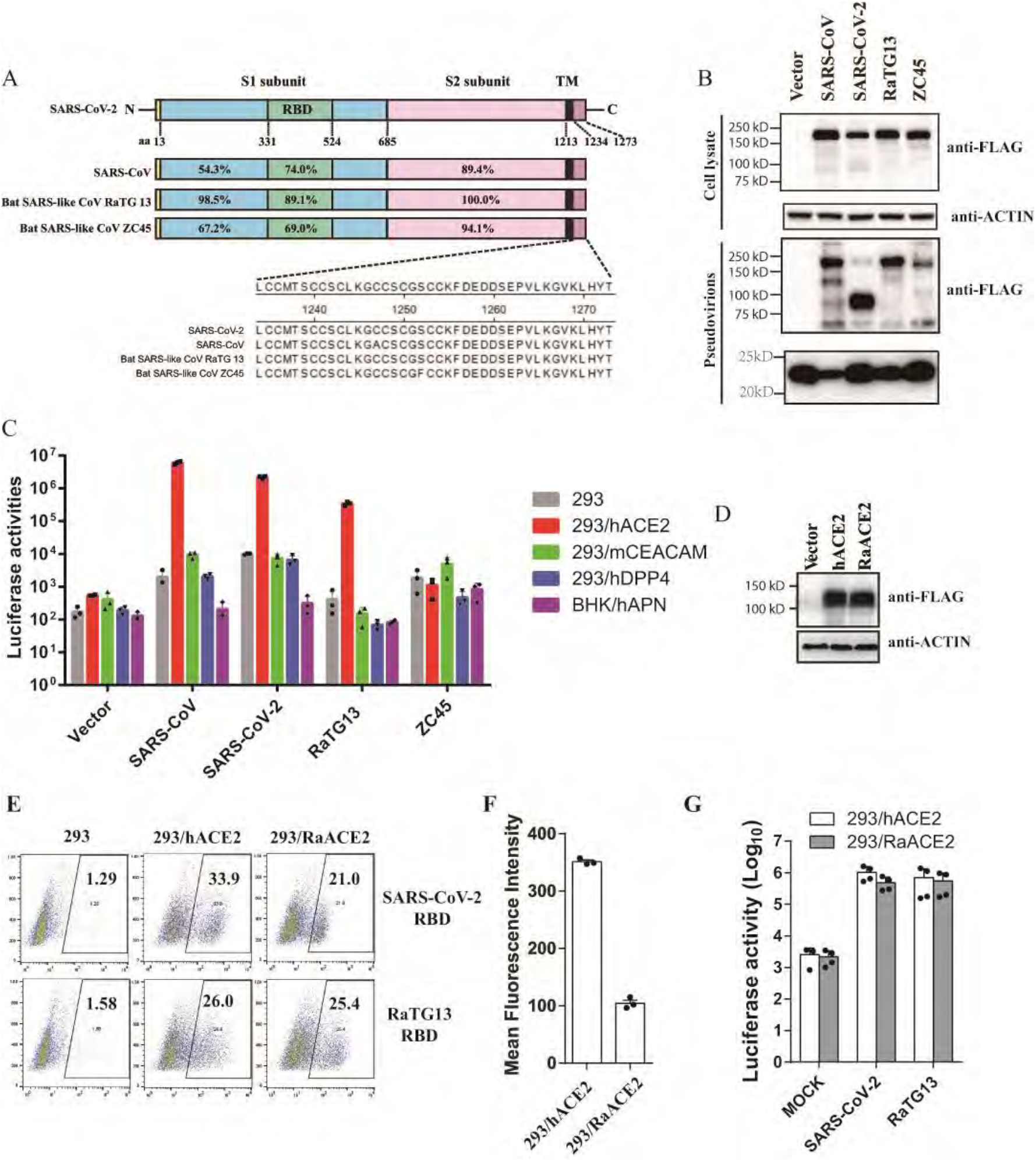
Bat SL-CoV RaTG13 uses hACE2 and RaACE2 for virus entry. (A) Schematic diagram of the full length of different CoV S proteins and the amino acid sequence identities of each region are shown in corresponding places. S1, receptor binding subunit; S2, membrane fusion subunit; TM, transmembrane domain. (B) Detection of the S proteins of SARS-CoV, SARS-CoV-2, Bat SL-CoV RaTG13 and ZC45 in cells lysates and pseudovirions by western blot. HEK293T cells transfected with either empty vector or plasmids encoding the indicated CoV S proteins were lysed at 40 hrs post transfection. The S proteins in cell lysates and pseudovirions were subjected to WB analysis by blotting with mouse monoclonal anti-FLAG M2 antibody. Actin and gag-p24 served as loading controls (cell lysate, top panel, pseudovirions, bottom panel). The full length S protein is about 180 kDa, while cleaved S protein is about 90 kDa. Experiments were done three times and the representative was shown. (C) Entry by RaTG13 S pseudovirons on different CoV receptors. Cells were spin-inoculated with indicated pseudovirions. At 48 hrs post inoculation, transduction efficiency was determined by measurement of luciferase activities. HEK293 cells (grey), HEK293/hACE2 (red), HEK293 cells stably expressing hACE2; 293/mCEACAM (green), HEK293 cells stably expressing mCEACAM, the MHV receptor; 293/hDPP4 (blue), HEK293 cells stably expressing hDPP4, the MERS-CoV receptor. BHK/hAPN(purple), BHK cells stably expressing hAPN, the hCoV-229E receptor; Experiments were done triplicate and repeated at least three times. One representative is shown with error bars indicate SEM. (D) Expression of Rhinolophus affinis ACE2 protein in HEK 293 cells. HEK 293 cells transiently transfected with the plasmids encoding either FLAG-tagged hACE2 or Rhinolophus affinis ACE2 (RaACE2) proteins were lysed at 40 hrs post-transfection. Expression of ACE2 proteins were detected by mouse monoclonal anti-FLAG M2 antibody. (E) Binding of hACE2 and RaACE2 by SARS-CoV-2 and RaTG13 RBDs. HEK 293 cells transiently expressing hACE2 or RaACE2 proteins were incubated with either SARS-CoV-2 RBD or RaTG13 RBD on ice, followed by rabbit anti-his tag antibodies and alexa-488 conjugated goat anti rabbit IgG, and analyzed by flow cytometry. The experiments were done three times, and one representative is shown. (F) Mean fluorescence intensities of the gated cells positive for SARS-CoV-2 RBD binding to 293/hACE2 and 293/RaACE2 cells in (E). (G) Entry of SARS-CoV, SARS-CoV-2, and RaTG13 S protein pseudovirions on 293/RaACE2 cells. Experiments were done three times, and one representative is shown with error bars indicating SEM. *P<0.05;**P<0.001 (compared with control by ANOVA followed by Dunnett’s multiple comparisons t test)

The level of S protein incorporation on pseudovirions was also evaluated. The S proteins of RaTG13 and ZC45 were efficiently incorporated into pseudovirions (Fig 1B and Supplementary Fig 1E). Next, we determined whether the RaTG13 and ZC45 S proteins can use any known coronavirus receptors for viral entry. The pseudovirions were used to transduce HEK293 cells stably expressing hACE2 (293/hACE2), HEK293 cells stably expressing hDPP4 (293/hDPP4), BHK cells stably expressing human aminopeptidase N (BHK/hAPN), or HEK293 cells stably expressing mouse carcinoembryonic antigen related cell adhesion molecule 1a (293/mCEACAM1a). SARS-CoV and SARS-CoV-2 S pseudovirions were used as controls. As expected, SARS-CoV and SARS-CoV-2 S pseudovirions utilized only hACE2, not mCEACAM1a, hDPP4, or hAPN, for virus entry (Fig 1C). While 293/hDPP4, BHK/hAPN, and 293/mCEACAM1a cells only showed background level of transduction with RaTG13 S pseudovirions, 293/hACE2 cells gave approximately 850-fold increase in luciferase activities over the HEK293 control when transduced by pseudovirions with RaTG13 S proteins, indicating that SL-CoV RaTG13 could use hACE2 as the entry receptor, in agreement with previous report^13,23^. In contrast, pseudovirions with ZC45 S protein did not transduce any cells effectively, indicating that SL-CoV ZC45 could not use any of them for virus entry.

Because RaTG13 virus was initially and only discovered in specimens from a single *Rhinolophus affinis* bat, we then investigated whether RaACE2 could also be the entry receptor for RaTG13 virus or not. The binding of RaACE2 to the S protein of RaTG13 was first evaluated. HEK293 cells transiently expressing RaACE2 proteins (Fig 1D) were incubated with soluble RaTG13 receptor binding domain (RBD) and their affinities were measured by flow cytometry. The RaTG13 RBD bound to RaACE2 proteins efficiently, at a level similar to that of hACE2 (Fig 1E, bottom panel). Of note, RaACE2 also bound to the SARS-CoV-2 RBD, but the affinity was significantly weaker than that of the hACE2/SARS-CoV-2 RBD (Fig 1E, top panel). The mean fluorescence intensity (MFI) of RaACE2/SARS-CoV-2 RBD interaction was less than 1/3 of that of the hACE/SARS-CoV-2 RBD (Fig 1F). RaTG13 RBD also demonstrated slightly weaker binding to hACE2 than SARS-CoV-2 RBD. Next, we determined whether RaACE2 could mediate the entry of RaTG13 and SARS-CoV-2 viruses. RaTG13 S pseudovions entered 293/RaACE2 cells at a level similar to hACE2, whereas SARS-CoV-2 S pseudovirions also transduced 293/RaACE2 cells efficiently, at slightly lower levels than hACE2 (Fig 1G). RaACE2 is a functional entry receptor for both RaTG13 and SARS-CoV-2 viruses.

Recently cats, civets, ferrets, minks, tigers, hamsters, dogs, and monkeys were reported to be susceptible to SARS-CoV-2 infection ^16-21^, and in silico analysis also showed that ACE2 from other animals might be able to mediate SARS-CoV-2 entry^22^. We next investigated which other animal ACE2 could confer susceptibility to RaTG13 virus entry. Sixteen different animal species (Table 1) were chosen, most of which are commonly found in wild animal meat markets in China, and we also included pangolins and two horseshoe bats (*Rhinolophus sinicus*), one from Yunnan (RS-YN bat), and the other from Hubei (RS-HB bat) in this study^27^, due to the discovery of some CoVs that are highly homologous to SARS-CoV and SARS-CoV-2 in them^9,28-30^. Among the 20 residues in hACE2 making direct contact with SARS-CoV-2 S proteins (Table 1), deer ACE2 differs in three positions with hACE2, squirrel has four residues that are different, ACE2s of fox, camel, pig, and RS-HB bat each have five residues that are different, RS-YN bat ACE2 has six residues that are different, ACE2s of pangolin, ferret, and guinea pig each have seven different, both rat and mouse ACE2s have eight residues different, and ACE2s of the remaining animals have nine or more residues different with hACE2 (Table 1). The plasmids encoding individual ACE2 proteins from these 16 different animal species (total 17, two horseshoe bats) were transfected into 293 cells, and the levels of their expression in 293 cells were determined by western blot (Fig 2A). While all ACE2 proteins were expressed in 293 cells (Fig 2A), expression levels varied among different ACE2 proteins, with the lowest for deer and snake ACE2s and the largest for hedgehog ACE2. The size for different ACE2 proteins also varied. While the deer ACE2 was the smallest, turtle ACE2 was the biggest. The deer ACE2 sequence we obtained from Genbank seems to lack the transmembrane domain (TMD) of ACE2, indicating that there might be different splicing variants of ACE2 in deer. We then investigated whether all different ACE2 proteins were present on the cell surface using a surface biotinylation assay. Except for the deer ACE2 protein, which lacked a TMD, most ACE2 proteins were present on the cell surface (Fig 2B). However, the levels of ACE2 proteins from guinea pigs and snakes were significantly lower on the cell surface than on the other surfaces. Integrin-β1 was used as the positive control for cell surface membrane proteins (Fig 2B). Because of the lack of TMD, the deer ACE2 was then removed from the rest of the analysis.

**Figure 2.**
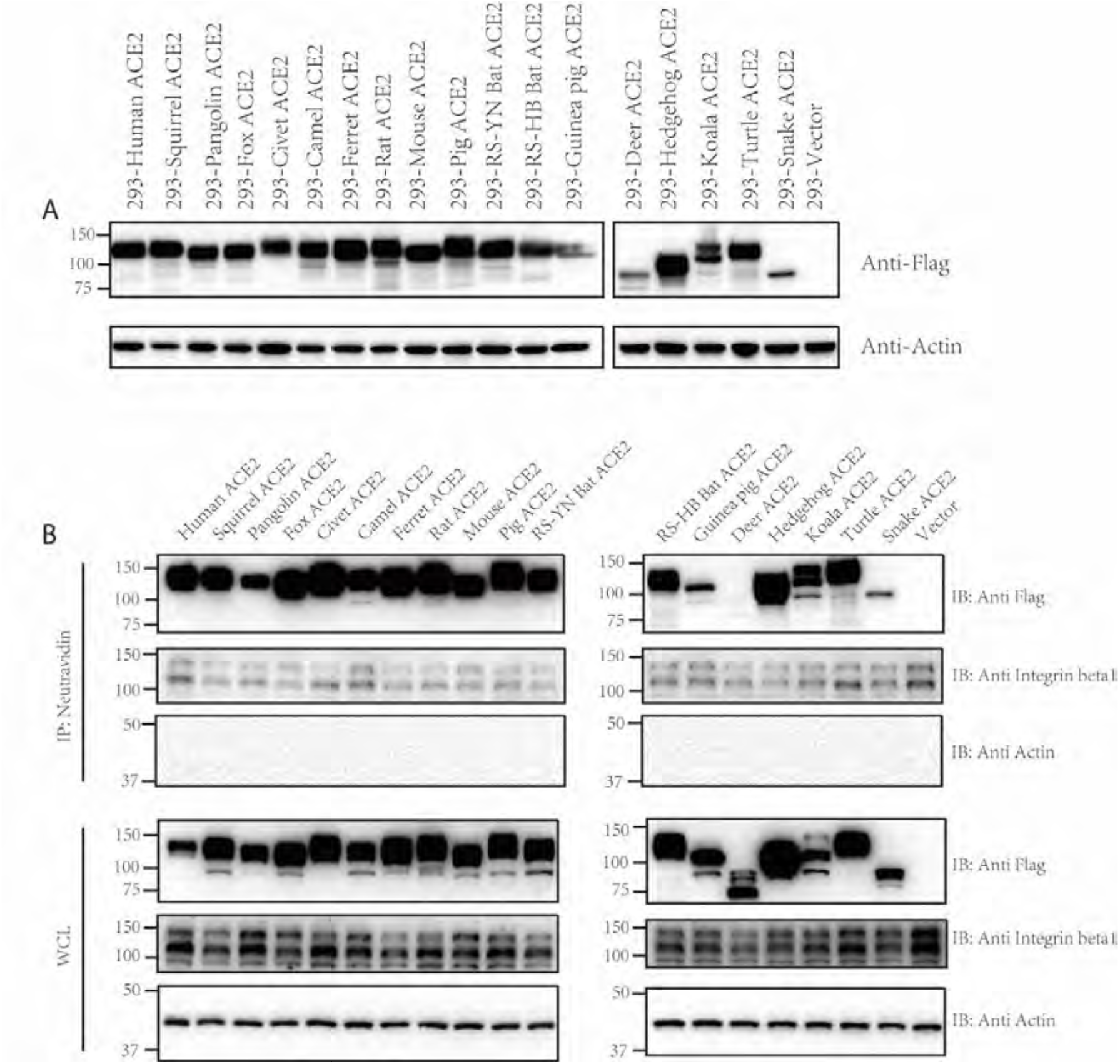
Expression and cell surface localization of ACE2 orthologs of various species. (A) Expression of ACE2s of different species in HEK293 cells. HEK293 cells were transfected with plasmids encoding FLAG tagged different ACE2s by PEI, and lyzed at 40 hrs post transfection. The expression of different ACE2 proteins in cell lysates was determined by western blotting using anti-FLAG M2 antibodies. The accession numbers for each ACE2 orthologs are as follows: human ACE2: NP_068576, squirrel ACE2: XP_026252505, pangolin ACE2: XP_017505746, fox ACE2: XP_025842512, civet ACE2: AAX63775, camel ACE2: XP_006194263, ferret ACE2: NP_001297119, rat ACE2: NP_001012006, mouse ACE2: NP_001123985, pig ACE2: NP_001116542, RS bat: AGZ48803, RS-HB bat: ADN93475, guinea pig ACE2: ACT66270, deer ACE2: XP_020768965, hedgehog ACE2: XP_004710002, koala ACE2: XP_020863153, turtle ACE2: XP_006122891, snake ACE2: ETE61880. (B) Analysis of different ACE2 proteins on cell surface by cell surface protein biotinylation assay. HEK293 cells transiently overexpressing different ACE2 proteins were labeled with EZ-link Sulfo-NHS-LC-LC-biotin on ice, and lysed with RIPA buffer. Biotinylated proteins were enriched with NeutrAvidin beads and analyzed by western blot using mouse monoclonal anti-FLAG M2 antibody. WCL, whole cell lysate.

Next we determined whether these different ACE2 proteins could bind to the S proteins of RaTG13. For comparison purposes, we also included SARS-CoV and SARS-CoV-2 in the rest of the experiments. HEK293 cells transiently expressing different ACE2 proteins were incubated with soluble RaTG13, SARS-CoV, and SARS-CoV-2 receptor binding domains (RBDs), and the percentage of cells that bound the RBD and the level of RBD bound to different ACE2 proteins were quantitated by flow cytometry (Fig 3 and Supplementary Fig 2). All RBDs of RaTG13, SARS-CoV-2, and SARS-CoV bound to HEK293 cells transiently expressing hACE2 protein, with RaTG13 RBD showing slightly lower levels of binding than SARS-CoV and SARS-CoV-2 RBDs (Supplementary Fig 2), consistent with the slightly lower transduction on 293/hACE2 by RaTG13 S pseudovirion than SARS-CoV and SARS-CoV-2 S pseudovirions (Fig 1C). Fox, camel, and pig ACE2 proteins also gave strong binding to RBDs of RaTG13, SARS-CoV, and SARS-CoV-2 at levels similar to hACE2 (Fig 3). In contrast, rat ACE2 proteins also bound to all three RBDs, but only at modest levels, ranging from 16% to 28% of hACE2. While squirrel and mouse ACE2 proteins bound strongly to RBDs of RaTG13 and SARS-CoV, they only bound to SARS-CoV-2 RBD at levels that were 36% and 12% of hACE2, respectively. In contrast, pangolin ACE2 proteins showed high affinity for both SARS-CoV and SARS-CoV-2 RBDs, but only weakly bound to RaTG13 RBD. SARS-CoV RBD bound civet and ferret ACE2 proteins at levels similar to hACE2, whereas SARS-CoV-2 RBD only showed binding to these ACE2 proteins at levels that were 24% and 15% of hACE2, respectively, and RaTG13 RBD only showed background level of binding. Of note, the RaTG13 and SARS-CoV-2 RBDs showed modest and strong binding to the ACE2 proteins of the RS-YN bat, respectively, but neither bound to the ACE2 proteins of the RS-HB bat (Fig 3A and 3B), in which there were seven S protein-interacting residues differing from those of RS-YN bat (Table 1). In contrast, SARS-CoV RBD showed modest but consistent binding to the RS-HB bat, not RS-YN bat (Fig 3C), reflecting the differences of receptor-contacting residues in RBDs among the three CoVs. None of ACE2 proteins from the other animal species showed any significant binding to either one of three RBDs. Overall, the fewer the number of critical binding residues that differ from hACE2 (Table 1), the higher the levels of binding detected. Both the RaTG13 and SARS-CoV-2 RBDs showed high affinity to ACE2 proteins of five different animals at levels of 60% or above that of hACE2, where SARS-CoV RBD bound to ACE2 proteins of seven different animals at 60% or above that of hACE2, indicating their potential broad range of hosts.

**Figure 3.**
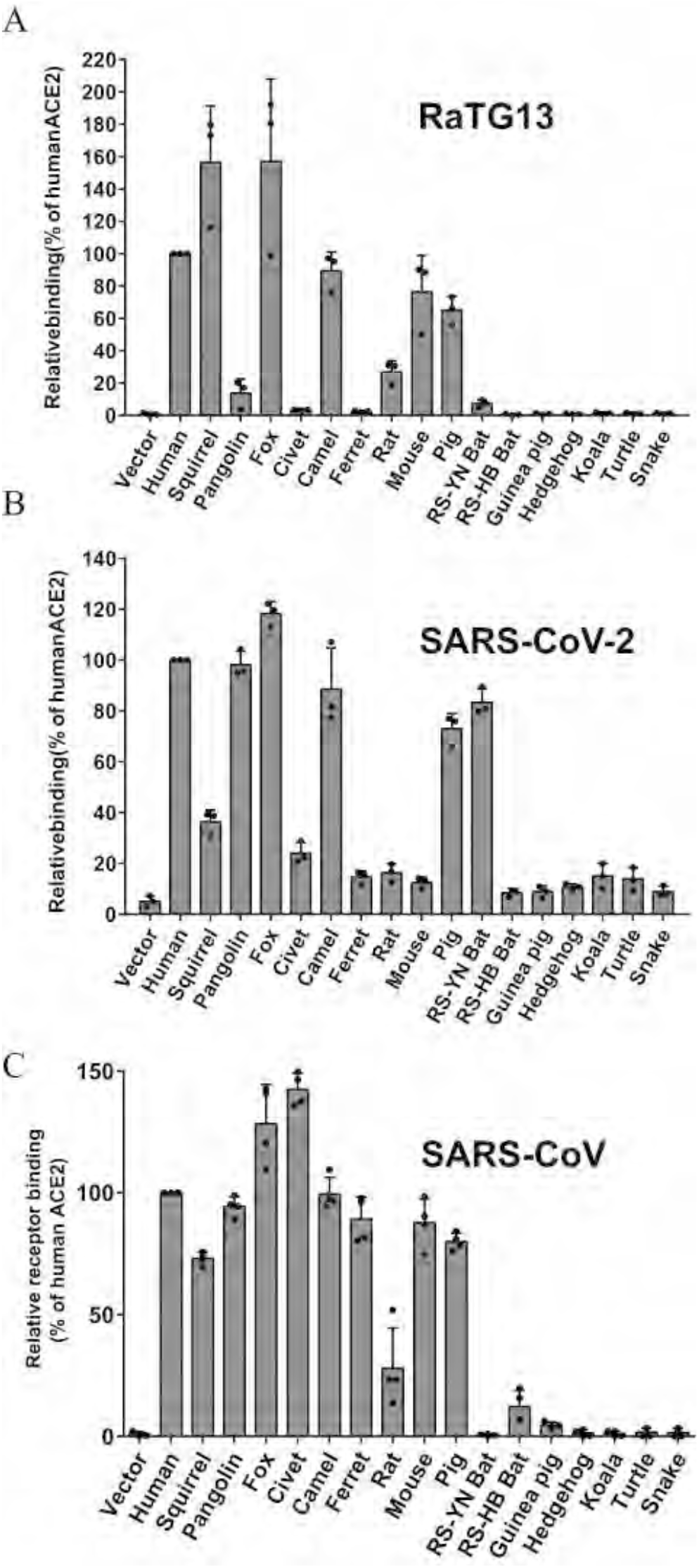
Binding of different ACE2 proteins by RBDs of bat SL-CoV RaTG13, SARS-CoV-2, and SARS-CoV. HEK293 cells transiently expressing different ACE2 cells were incubated with either RaTG13 (A), SARS-CoV-2 (B), or SARS-CoV (C) RBDs, followed by rabbit anti-His tag antibodies and Alexa-488 conjugated goat anti rabbit IgG, and analyzed by flow cytometry. The experiments were done at least three times. The results of percentage of positive cells from hACE2 binding were set to 100%, the rest was calculated as percentage of hACE2 binding according to results in flow cytometry analysis. Data are shown as the means ± standard deviations.

Membrane fusion is a prerequisite step for virus entry. We next evaluated the effect of different animal ACE2 proteins on the S protein of RaTG13 mediated membrane fusion by cell-cell fusion assay. SARS-CoV and SARS-CoV-2 S proteins were also used for comparisons. In agreement with our previous report^10^, HEK293 cells transiently expressing hACE2 proteins showed extensive syncytium formation when coincubated with 293T cells overexpressing eGFP and SARS-CoV or SARS-CoV-2 S proteins in the presence of trypsin (Fig 4A). Syncytia were also induced when RaTG13 S protein expressing cells were overlaid on HEK293 cells transiently expressing hACE2 with trypsin. However, the frequency and size of the syncytium were much lower and smaller than the S proteins of SARS-CoV and SARS-CoV-2 (8.7% for RaTG13, 37.3% for SARS-CoV-2, and 29.1% for SARS-CoV) (Fig 4A, 4B, 4C, 4D and Supplementary Figure 3). Of note, HEK293 cells expressing fox and rat ACE2 proteins, and to a lesser extent, squirrel and mouse ACE2 proteins showed significantly higher amount of syncytium formation than hACE2 when mixed with RaTG13 S protein expressing cells and trypsin (Fig 4B), although all were present on the cell surface at similar level (Fig 2B). Camel ACE2 also induced syncytia at a level similar to hACE2, whereas civet, ferret and pig ACE2s showed syncytia at 65%, 49% and 61% of hACE2, respectively. None of the other animal ACE2s, including ACE2s from two horseshoe bats, induced marked syncytium formation by RaTG13 S proteins. Overall, the cell-cell fusion results were largely in agreement with the ability of ACE2 proteins binding RaTG13 RBD.

**Figure 4.**
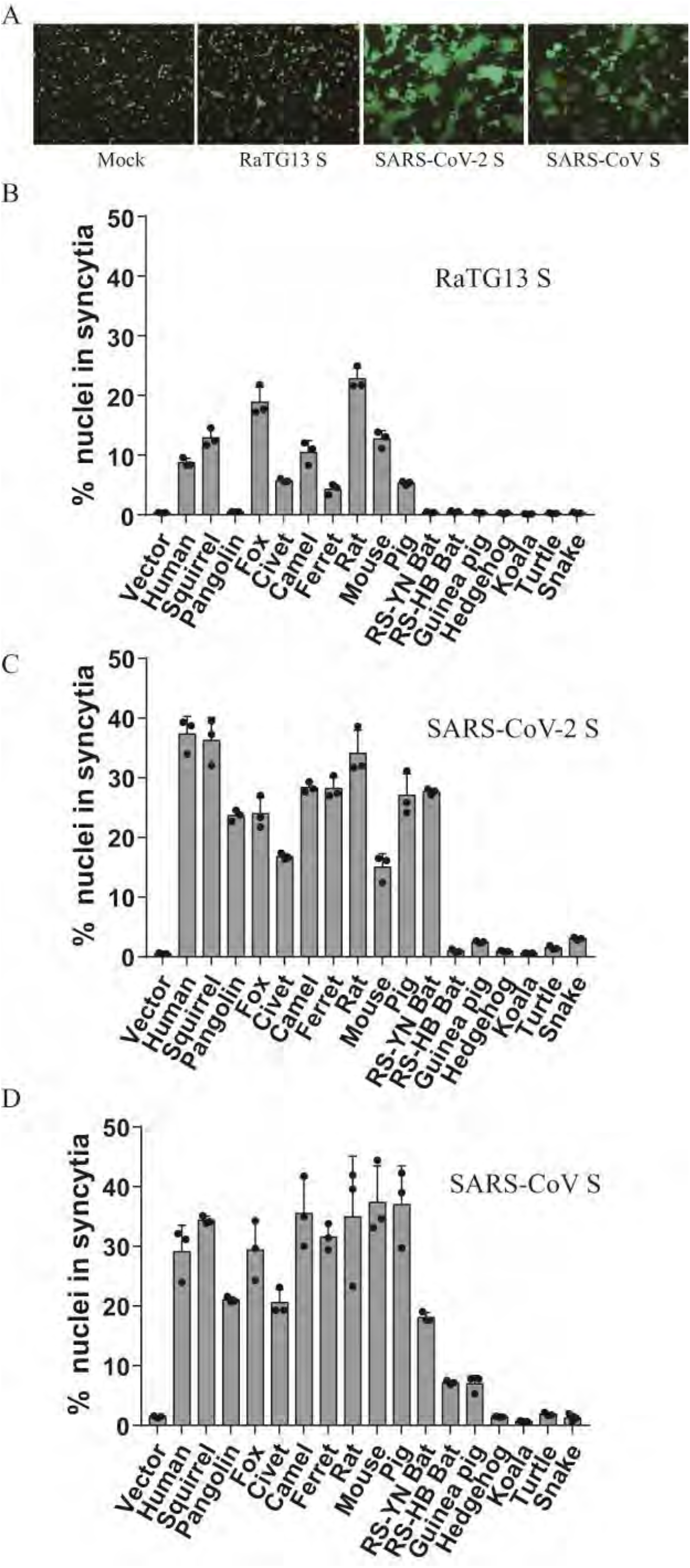
Cell–cell fusion mediated by RaTG13, SARS-CoV, and SARS-CoV-2 spike proteins. HEK293T cells transiently expressing eGFP and spike proteins of either RaTG13, SARS-CoV, or SARS-CoV-2 were detached with trypsin, and overlaid on different ACE2 expressing HEK293 cells. After 4 hrs of incubation, images were taken. (A) Representative images of syncytia for hACE2; (B-D) Percentage of nuclei in syncytia induced by RaTG13 S (B), SARS-CoV-2 S (C), and SARS-CoV S (D). Syncytium formation for each image was quantified by counting the total nuclei in syncytia and total nuclei in the image and calculated as the percentage of nuclei in syncytia, and three images were selected for each sample. Experiments were done three times, and one representative is shown with error bars indicating SEM. The scale bar indicates 250 μm.

The S protein of SARS-CoV-2 induced extensive syncytia on HEK293 cells transiently expressing squirrel, pangolin, fox, civet, camel, ferret, rat, mouse, pig, and RS-YN bat ACE2 proteins (Fig 4C), although ferret, rat, and mouse ACE2 protein only showed binding to the SARS-CoV-2 RBD slightly above background level (Fig 3B). Because several recent studies reported that mouse ACE2 is not susceptible to SARS-CoV-2 infection^3,31^, we repeated the cell-cell fusion experiments multiple times with mouse ACE2 plasmids, prepared with extra caution and verified by sequencing, and significant amount of syncytia were still detected (Supplementary Figure 4). All other animal ACE2 proteins gave background levels of syncytia, consistent with their inability to bind to the SARS-CoV-2 RBD. The overall pattern of SARS-CoV S protein mediated syncytium formation on different animal ACE2 expressing 293 cells was similar to that of SARS-CoV-2 S protein. Of note, although RS-YN bat ACE2 did not show any marked binding to SARS-CoV RBD, it induced SARS-CoV S protein mediated syncytium formation at a level of 62% of hACE2. HEK293 cells expressing RS-HB bat and guinea pig ACE2 also induced noticeable syncytium formation upon addition of SARS-CoV S expressing 293T cells and trypsin (Fig 4D). Overall, the S proteins of SARS-CoV and SARS-CoV-2 showed much higher fusogenicity than the RaTG13 S proteins.

Next we investigated whether ACE2 proteins from different animal species could mediate virus entry by the RaTG13, SARS-CoV, and SARS-CoV-2 S proteins. Lentiviral pseudovirions with VSV-G protein were used as a positive control. As expected, all cells were susceptible to VSV-G pseudoviron transduction (Fig 5). Compared to vector control, 293/hACE2 cells showed an over 3500-fold increase in luciferase activities when transduced with RaTG13 S pseudovirions (Fig 5A). Over a 1000-fold increase of transduction was also detected in HEK293 cells transiently overexpressing squirrel, fox, camel, and mouse ACE2 proteins (Fig 5A), indicating that they might be susceptible to RaTG13 infection. Rats and pigs also seem to be susceptible to RaTG13 infection, and their ACE2s resulted in over 225- and 630-fold increases in luciferase activities (Fig 5A), respectively, when transduced by RaTG13 S pseudovirions, largely in agreement with their ability to bind to RaTG13 RBD. In contrast, although civet and ferret ACE2 only showed minimal binding to the RaTG13 RBD, they gave close to 500- and 90-fold increases in luciferase over the vector control (Fig 5A), respectively, when transduced by RaTG13 pseudovirons, indicating that they might also be susceptible to RaTG13 infection. Of note, neither horseshoe bat ACE2s t showed high susceptibility to RaTG13 S pseudovirion transduction. While RS-YN bat ACE2 gave an approximately 13-fold increase in transduction over the vector control, RS-HB bat ACE2 only showed a background level of transduction (Fig 5A). None of the other animal ACE2s showed a significant increase in virus entry by the RaTG13 S protein.

**Figure 5.**
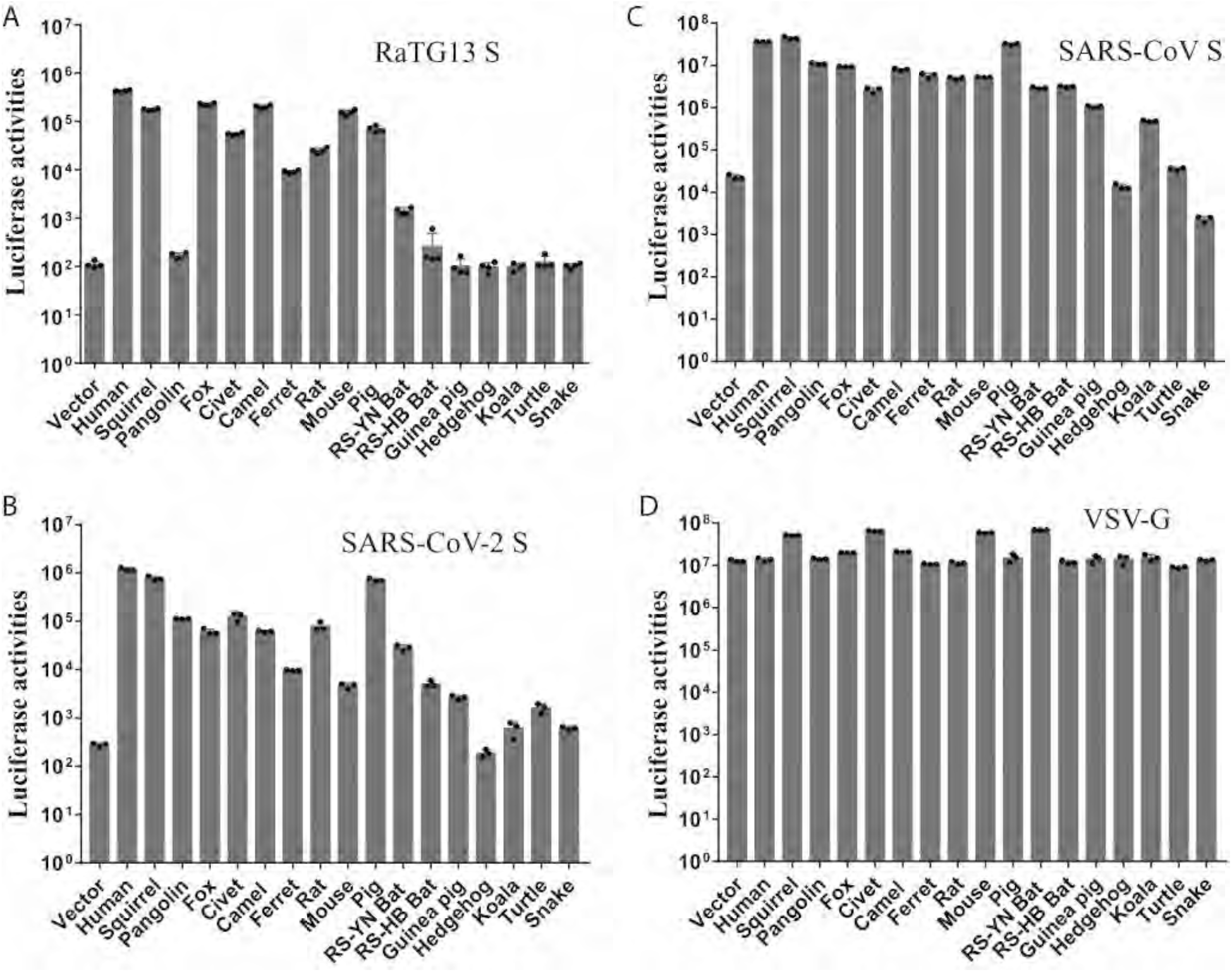
Entry mediated by the S protein of RaTG13, SARS-CoV-2, and SARS-CoV on cells expressing different ACE2 proteins. HEK-293 cells transiently expressing different ACE2 proteins were transduced with RaTG13 S pseudovirions (A), SARS-CoV-2 S pseudovirions (B), SARS-CoV S pseudovirions (C), and VSV-G pseudovirions (D). Experiments were done in triplicate and repeated at least three times. One representative is shown with error bars indicating SEM.

Overall, SARS-CoV-2 S protein pseudovirions showed similar levels of host ranges to RaTG13 among the different ACE2s we tested (Fig 5B). However, they differed dramatically in susceptibility to pangolins and mice. Pangolin ACE2 was susceptible to SARS-CoV-2 S-mediated transduction (Fig 5B), but not RaTG13 (Fig 5A), whereas mouse ACE2 was fully susceptible to RaTG13 transduction (Fig 5A), but limited to SARS-CoV-2 (Fig 5B). SARS-CoV-2 S pseudovirions showed only 0.4% of hACE2 transduction in mouse ACE2 (Fig 5C). HEK293 transiently overexpressing squirrel and pig ACE2s gave a level of transduction similar to that of hACE2 (Fig 5B), although squirrel ACE2 bound to the SARS-CoV-2 RBD at a level of less than 40% of that of hACE2 (Fig 3B). Fox, civet, camel, rat, or RS-YN bat ACE2 proteins also gave an over 100-fold increase in luciferase activities (Fig 5C), indicating that these animals might be susceptible to SARS-CoV-2 infection. Ferret ACE2 also showed an approximately 35-fold increase in transduction (Fig 5C), in agreement with recent studies showing that ferrets were susceptible to SARS-CoV-2 infection^32,33^. RS-HB bat and guinea pig ACE2 proteins only gave approximately 15- and 10-fold increases in transduction by SARS-CoV-2 S pseudovirions, in agreement with their low affinity of binding to SARS-CoV-2 S protein. Compared to RaTG13 and SARS-CoV-2, SARS-CoV S showed broader host range. HEK293 cells transiently expressing squirrel, pangolin, fox, civet, camel, ferret, rat, mouse, pig, RS-YN bat, RS-HB bat, guinea pig, and koala ACE2 showed marked increases in luciferase activities (Fig 5B), when transduced by SARS-CoV S pseudovirions. Of note, both RS-YN and RS-HB bat ACE2 proteins showed a more than 100-fold increase in transduction (Fig 5C), compared to vector control, although they exhibited substantial differences in binding to the SARS-CoV RBD and syncytium formation (Fig 4D).

To identify the residues in the S proteins of RaTG13 and SARS-CoV-2 critical for the interaction and recognition of ACE2 in different animal species, we applied in silico analyses of SARS-CoV-2/RaTG13 RBDs and different ACE2 interactions using HAWKDOCK and PYMOL (Supplementary Fig 5), particularly focusing on mouse and pangolin ACE2. In the SARS-CoV-2 RBD/hACE2 crystal structure, interactions between hACE2 and the SARS-CoV-2 RBD complex consist of extensive network of hydrogen bonding salt bridges and hydrophobic interactions (Supplementary Fig 5A, 5B, 5C and 5D)^12,14^. F28, L79, M82, and Y83 in hACE2 form a hydrophobic pocket interacting with the critical F486 in the SARS-CoV-2 S protein^34^ (Supplementary Fig 5B). L79T, M82S, and Y83F changes in mouse ACE2 might collapse this hydrophobic pocket and weaken the interaction with F486 in S protein (Supplementary Fig 5E). The D30N change in mouse ACE2 likely abrogates the salt bridge with K417 in the S protein of SARS-CoV-2, and K31N and K353H changes in mouse ACE2 also likely disrupt the hydrogen bonding network between ACE2 and SARS-CoV-2 S protein, resulting in mouse ACE2 acting as a poor receptor for SARS-CoV-2. In contrast, K439, Y493 and Y498 in the RaTG13 S protein might make hydrogen bonds with Q325, N31 and Q42 in mouse ACE2(Supplementary Fig 5E), resulting in an increase in the overall affinity between mouse ACE2 and the RaTG13 S protein (Fig 3A) and virus entry by the RaTG13 virus (Fig 5A).

Pangolin ACE2 differs from human ACE2 at seven critical positions making contact with the RBD (Table 1), of which three (E30, E38, and I79) are homologous and four (E24, S34S, N82, and H354) are different. While these changes do not affect SARS-CoV-2 RBD binding to pangolin ACE2 (Supplementary Fig 5F), they appear to be detrimental to RaTG13 RBD binding (Fig 3A) and virus entry (Fig 5A). In silico analysis showed that Y449F, E484T and Q493Y changes in RaTG13 S protein might disrupt their hydrogen bonding with E38, K31, and E35 of pangolin ACE2, respectively (Supplementary Fig 5F), resulting in weak interaction between RaTG13 S protein and pangolin ACE2 and poor transduction efficiency of pangolin ACE2 by RaTG13 S protein (Fig 5A).

Based on the results from the in silico analysis, we selected residues 449, 484, 493, and 498 in the S proteins for further studies (Fig 6A). Single mutations F449Y, T484E, Y493Q, and Y498Q were introduced into the RaTG13 S protein, and individual Y449F, E484T, Q493Y, and Q498Y mutations were also introduced into the SARS-CoV-2 S protein. All mutant RaTG13 S proteins were expressed as well as WT in HEK293T cells and incorporated into pseudovirion efficiently (Fig 6B), whereas all mutant SARS-CoV-2 S proteins except for Y493Q were expressed and incorporated into pseudovirions at levels similar to WT (Fig 6C). Because Q493Y mutation in SARS-CoV-2 S had significant effect on S protein incorporation into pseudovirions, they were removed from further analysis. We then determined whether any mutations affected virus entry using hACE2. While both the F449Y and Y498Q mutations in the RaTG13 S protein significantly reduced virus entry into 293/hACE2 cells (Fig 6D), indicating that both F449 and Y498 of RaTG13 might be critical for virus entry through hACE2, the Y493Q substitution in the RaTG13 S protein significantly increased transduction into hACE2 cells, suggesting that Q might be advantageous at position 493 for interaction with hACE2. In contrast, only the Y449F mutation in the SARS-CoV-2 protein showed greater than 50% reduction in infectivity in 293/hACE2 cells (Fig 6E). Next, we investigated whether any mutations influenced virus entry into mouse and pangolin ACE2s. The overall patterns of mutant RaTG13 S pseudovirion infectivity in mouse ACE2 cells were very similar to those on hACE2 cells except for T484E (Fig 6F and 6G). The F449Y, T484E, and Y498Q mutant RaTG13 S proteins showed a significant reduction in infectivity on mouse ACE2, whereas the Y493Q substitution slightly increased transduction on mouse ACE2 by RaTG13 S pseudovirions (Fig 6F). These results suggested that residues 449, 484, and 498 of RaTG13 S protein might also be important for interaction with mouse ACE2 and that Q might be preferred over Y at position 493 of the RaTG13 S protein to interact mouse ACE2. None of the mutations could significantly rescue the infection of RaTG13 S pseudovirions on pangolin ACE2 expressing cells (Fig 6F). The effects of individual mutations in SARS-CoV-2 S proteins on virus entry through pangolin ACE2 were relatively limited (Fig 6G), and very similar to those in hACE2 cells (Fig 6E). Y449F and E484T mutants showed slightly over 50% and 30% reduction in infectivity in pangolin ACE2-expressing cells, respectively, whereas Q498Y mutations in SARS-CoV-2 had no effect on virus entry into pangolin ACE2 expressing cells. Strikingly, mutant E484T and Q498Y SARS-CoV-2 S proteins increased transduction on mouse ACE2 expressing cells by more than 16 and 70-fold, respectively, indicating that residues 484 and 498 of SARS-CoV-2 S proteins might play critical roles in determining receptor usage of mouse ACE2.

**Figure 6.**
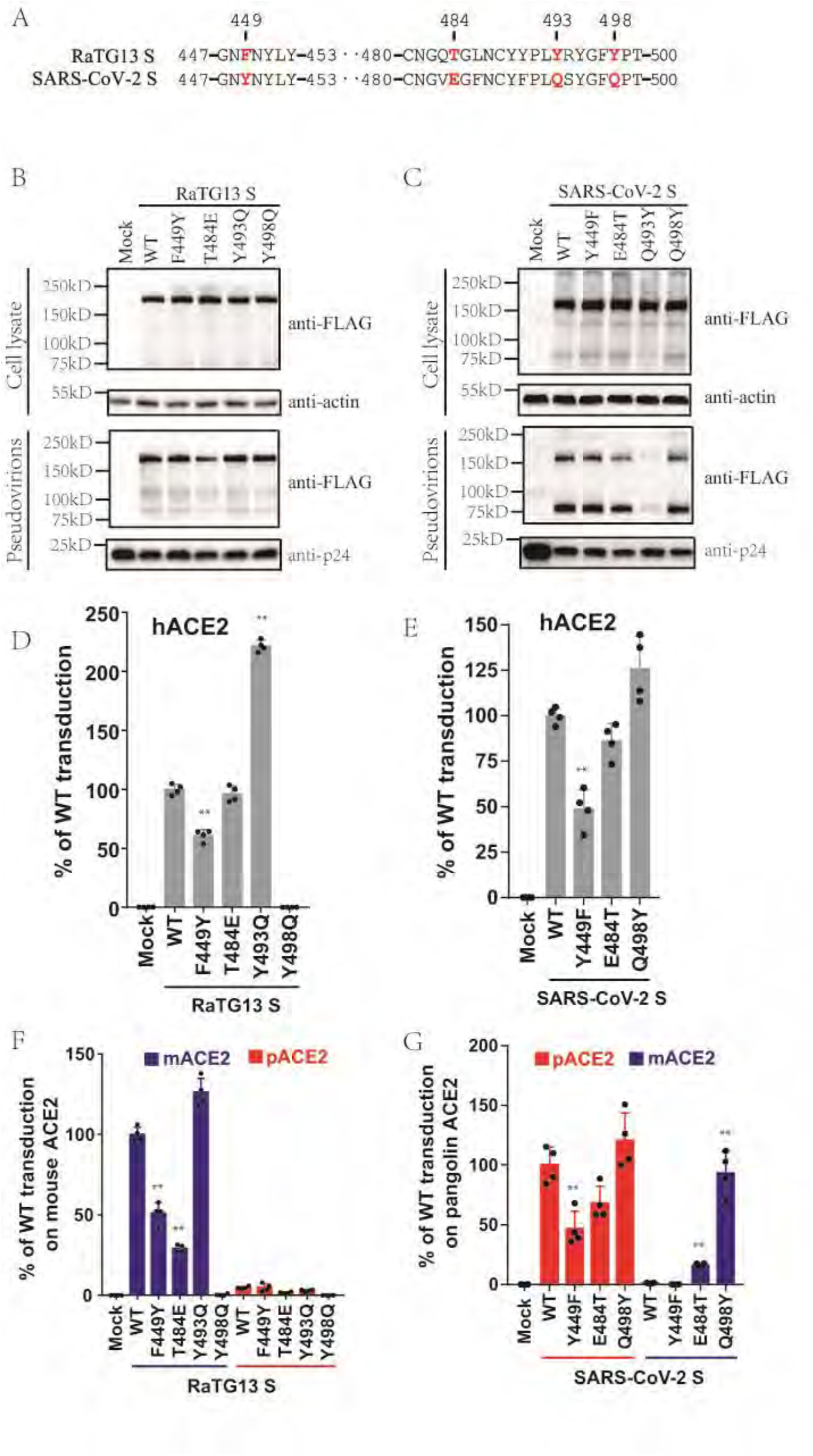
Entry of lentiviral pseudovirions with mutant RaTG13 S and SARS-CoV-2 S proteins on 293/hACE2, 293/mouse ACE2, and 293/pangolin ACE2 cells. (A) Alignment of partial amino acid sequences of RaTG13 and SARS-CoV-2 S proteins. Residues 449, 484, 493, and 498 are labeled in red. Detection of mutant S proteins in cells lysates and pseudovirions by western blotting using a mouse monoclonal anti-FLAG M2 antibody. (B) RaTG13 S. (C) SARS-CoV-2 S. Top panel, cell lysate; bottom panel, pseudovirions; β-actin and HIV p24 were used as loading controls. (D)(E) Entry of pseudovirons with mutant RaTG13 (D) and SARS-CoV-2 (E) S proteins on 293/hACE2 cells. Pseudovirions carrying mutant S proteins were inoculated on 293/hACE2 cells. After 40 hrs incubation, transduction efficiency was determined by measuring the luciferase activities in cell lysate. Transduction from WT pseudovirions was set as 100%. Experiments were done in quadruplicate and repeated at least three times, and one representative was shown with SEM. (F) Entry of pseudovirons with mutant RaTG13 S proteins on 293 cells expressing mouse (blue) and pangolin (red) ACE2 proteins. Transduction from WT pseudovirions on mouse ACE2 cells was set as 100%. (G) Entry of pseudovirons with mutant SARS-CoV-2 S proteins on 293 cells expressing mouse (blue) and pangolin (red) ACE2 proteins. Transduction from WT pseudovirions on pangolin ACE2 cells was set as 100%. The experiments were performed in quadruplicate with at least three replications and the representative data are shown with SEM. *P<0.05;**P<0.001 (compared with respective WT control by ANOVA followed by Dunnett’s multiple comparisons t test)

## Discussion

Viral entry is the first step for zoonotic transmission, and the interaction between the host receptor and viral S protein determines the host range and viral tropism. Although the origin of SARS-CoV-2 remains unknown, RaTG13 has been speculated to be the possible origin of SARS-CoV-2^3,8,9,35^, because the genomes of SARS-CoV-2 and bat SL-CoV RaTG13 share the highest nucleotide sequence identity. Here we showed that, although both SARS-CoV-2 and RaTG13 could use hACE2 for virus entry, the S proteins of the two CoVs have marked differences in biological properties, in terms of their affinity to ACE2s of different animal species, their fusogenicity in membrane fusion, and virus entry using different ACE2 proteins, especially for pangolin and mouse ACE2s, supporting the hyothesis that SARS-CoV-2 might not arise from RaTG13 virus directly, consistent with previous analysis^36,37^.

The RaTG13 virus was originally found in specimens from a *Rhinolophus affinis* bat^3^, indicating that the *Rhinolophus affinis* bat might be a natural host for the RaTG13 virus. Our finding of RaACE2 as a functional entry receptor for RaTG13 virus (Fig 1E and 1G) provides the first direct evidence of this. In fact, RaACE2 was almost as efficient as hACE2 in binding to RaTG13 RBD and facilitating entry of RaTG13 S pseudovirions (Fig 1E and 1G). In contrast, SARS-CoV-2 clearly favored hACE over RaACE2 for receptor binding and modestly favor hACE2 over RaACE2 for virus entry (Fig 1E, 1F, and 1G), reflecting possible adaptation of SARS-CoV-2 in human beings or an unknown intermediate host, if SARS-CoV-2 evolved from RaTG13 or a RaTG13-like virus. There are four residues (R24, I27, N31, and N82) in RaACE2 that differ from hACE2 (Table 1). In silico analysis revealed that K31N and M82N changes in RaACE2 likely reduce hydrogen and hydrophobic interactions with the SARS-CoV RBD (Supplementary Fig 6), respectively, resulting in a decrease in overall affinity. In contrast, R24 in RaACE2 likely forms an extra hydrogen bond with S477 in the RaTG13 RBD but not S477 in SARS-CoV-2 (distance:2.8 Å vs 4.1 Å), stabilizing the interaction between RaACE2 and the RaTG13 RBD.

Identification of a direct natural animal reservoir and/or zoonotic intermediate host of SARS-CoV-2 is essential to prevent future emergence and re-emergence of SARS-CoV-2 or SARS-CoV-2 like viruses. Recently, several novel pangolin CoVs were discovered in Malayan pangolins rescued during an anti-smuggling campaign in Guangdong, China^9,28-30^. Among them, one RBD was almost identical to the SARS-CoV-2 RBD in terms of amino acid sequence, except for one single noncritical residue^9^, leading to the hypothesis that SARS-CoV-2 might result from recombination of RaTG13-like CoV and pangolin CoV and pangolin might be the intermediate host for SARS-CoV-2^9,28,30^. Pangolin ACE2 not only showed strong binding to SARS-CoV-2 S protein (Fig 3B) and triggered large syncytia mediated by SARS-CoV-2 S protein (Fig 4C), but HEK293 cells expressing pangolin ACE2 were also highly susceptible to SARS-CoV-2 S protein mediated virus entry, suggesting that pangolin should be susceptible to SARS-CoV-2 infection. However, RaTG13 RBD only showed very limited binding to pangolin ACE2 (Fig 3B), its S protein only induced background level of syncytia on HEK293 cells transiently expressing pangolin ACE2 (Fig 4C), and RaTG13 S pseudovirions also only gave background level of transduction on HEK293 cells transiently expressing pangolin ACE2 (Fig 5A), indicating that RaTG13 virus might not be able to infect pangolin. This raises the question of whether pangolins could be intermediate hosts for SARS-CoV-2 if RaTG13 or RaTG13-like viruses could not infect pangolins. Moreover, pangolins are solitary animals, and infection by these pangolin CoVs is lethal for most pangolins^9^, suggesting that these pangolin CoVs might not be native to pangolins. Recent studies on 334 *Sunda pangolins* did not find any CoVs or other potential zoonotic viruses in these animals^38^, further supporting that pangolins might not be reservoir hosts for these pangolin CoVs. Where, when, and how these pangolins acquired these CoVs remain elusive.

Among the 17 different ACE2s we tested, squirrel and pig ACE2s were highly susceptible to transduction by all SARS-CoV, SARS-CoV-2, and RaTG13 S pseudovirions (Fig 5), although recent studies reported that pigs might not be permissive to SARS-CoV-2 infection^17,39^, likely resulting from low level of expression of ACE2 proteins on pig respiratory track^22^. Fox, civet, camel, ferret, and rat were also susceptible to virus entry by all three S pseudovirions (Fig 5), indicating the potential broad host range of all three viruses. Ferret has been used as a SARS-CoV-2 infection and transmission model^17,32,33,40^. Whether any of these other susceptible animals could be used as animal models for SARS-CoV-2 remains to be determined, especially for rats, which are cheaper and widely available. More importantly, whether any of these susceptible animals might be potential intermediate hosts for SARS-CoV-2 warrants further investigation.

Of note, while SARS-CoV-2 could bind and use pangolin ACE2 for virus entry, its ability to use mouse ACE2 was very limited, and conversely, RaTG13 could bind and use mouse ACE2 for virus entry, but not pangolin ACE2. Among the 20 residues making direct contact with SARS-CoV-2 S proteins (Table 1), mouse ACE2 protein differs at eight RBD-interacting residues from human ACE2 (Table 1), whereas pangolin ACE2 has seven critical positions differing from hACE2 (Table 1). In silico analyses showed that multiple amino acid changes in mouse ACE2, including D30N, K31N, and K353H, likely disrupt the salt bridge and hydrogen bonding network between ACE2 and SARS-CoV-2 S protein, whereas several changes in RaTG13 S protein, including N439K, F486L, Q493Y, and Q498Y (N, F, Q, Q from SARS-CoV-2 S and K, L, Y, Y from RaTG13 S), might reestablish the interactions with mouse ACE2 (Supplementary Fig 5E). This notion is strongly supported by the results for the Q498Y mutation in the SARS-CoV-2 S protein and the Y498Q mutation in the RaTG13 S protein. Replacement of Q498 with Y increased the infectivity of SARS-CoV-2 S pseudovirions on mouse ACE2 expressing cells by more than 70-fold. In contrast, substitution of Y498 with Q almost abrogated transduction by RaTG13 S pseudovirions on mouse ACE2, indicating the importance of residue 498 of both the RaTG13 and SARS-CoV-2 S proteins in the recognition of the mouse ACE2 protein. Of note, Q498 mutations were found in two recent mouse adapted SARS-CoV-2 strains, Q498H in one^41^, and Q498T in the other^31^. We did not identify any residue in the S protein essential for interacting with pangolin ACE2. Y449F, E484T, and Q498Y substitutions in the SARS-CoV-2 S protein had moderate effect on virus entry into pangolin ACE2 cells, and none of the mutations in the RaTG13 S protein significantly increased virus infectivity in 293/pangolin ACE2 cells.

RS-YN bat ACE2 showed strong binding to SARS-CoV-2, induced syncytium formation effectively, and was susceptible to transduction by SARS-CoV-2 S pseudovirions, consistent with previous reports^3^. In silico analysis (Supplementary Figure 5) revealed that Y449, E484, and Q493 in SARS-CoV-2 S could form hydrogen bonds with D38, K31, and T34 in RS-YN bat ACE2, resulting in strong binding of SARS-CoV-2 RBD with RS-YN bat ACE2. In contrast, although RS-HB bat ACE2 has only a 5 amino acid difference from hACE2 and one fewer than RS-YN bat ACE2, it only showed a background level of binding to SARS-CoV-2. K31E, T34S, and D38N changes in RS-HB bat ACE2 might disrupt their hydrogen bonding with E484, Q493, and Y449 of SARS-CoV-2 RBD, respectively, critical for SARS-CoV-2 RBD and bat ACE2 interaction (Supplementary Figure 5). Both RS bat ACE2s seem to be poor receptors for RaTG13 virus. Y449F, E484T, and Q493Y changes in RaTG13 might abolish those critical hydrogen bonds, leading to very limited binding to both RS bat ACE2 (Fig 3B), background level of syncytium formation (Fig 4B), and limited virus entry by RaTG13 S pseudovirions (Fig 5A). Whether failure of SARS-CoV-2and RaTG13 using RS-HB bat ACE2 for virus entry might result from pathogen-driven host revolution remains to be determined. Although SARS-CoV-2 likely evolved from bat-CoV RaTG13 or RaTG13-like bat-CoV with or without recombination with other CoVs, the difference in susceptibility of the two RS bat ACE2s between SARS-CoV-2 and RaTG13 raises two important questions: 1. Which bat species other than *Rhinolophus affinis* might harbor RaTG13 or RaTG13-like virus? 2. How does RaTG13 or RaTG13-like CoV evolve to SARS-CoV-2?

In summary, we determined the susceptibility of bat-CoV RaTG13 to 17 diverse animal ACE2s and compared them with those of SARS-CoV-2 and SARS-CoV. We found that RaACE2 is an entry receptor for RaTG13 and SARS-CoV-2. All three CoVs likely have a broad host range with SARS-CoV being the broadest, and mice, not pangolins, are susceptible to RaTG13 infection, whereas pangolins, not mice, are susceptible to SARS-CoV-2 infection. Residues 484 and 498 in the S protein play critical roles in the recognition of mouse and human ACE2.

## Materials and Methods

### Constructs and plasmids

Codon-optimized cDNA (sequences are shown in Supplementary Table 1) encoding SARS-CoV-2 S protein (QHU36824.1),SARS-CoV S protein (AAP13441.1) and S proteins of SARS-like bat CoV RaTG13 (MN996532.1) and ZC45 lacking C-terminal 19 amino acids (aa)were synthesized and cloned into the eukaryotic cell expression vector pCMV14-3×Flag between the *Hind III* and *Xba I* sites. The VSV-G encoding plasmid and lentiviral packaging plasmid psPAX2 were obtained from Addgene (Cambridge, MA). The pLenti-GFP lentiviral reporter plasmid that expresses GFP and luciferase was generously gifted by Fang Li, Duke University. The cDNAs encoding ACE2 orthologs (Table 1) were synthesized by Sango Biotech (Shanghai, China) and cloned into the pCMV14-3×Flag vector between the *Hind III* and *BamH I* sites. All the constructs were verified by sequencing.

### Cell lines

Human embryonic kidney cell lines 293 (#CRL-1573) and 293T expressing the SV40 T-antigen (#CRL-3216) were obtained from ATCC (Manassas, VA, USA), HEK239 cells stably expressing recombinant human ACE2 (293/hACE2), baby hamster kidney fibroblasts stably expressing recombinant human APN (BHK/hAPN), HEK239 cells stably expressing recombinant human DPP4 (293/hDPP4), HEK-293 cells stably expressing murine CEACAM1a (293/mCEACAM1a)were established in our lab. All above cells were maintained in Dulbecco’s MEM containing 10% fetal bovine serum (FBS) and 100 units of penicillin, 100 μg of streptomycin, and 0.25 μg of fungizone (1% PSF, Gibco) per milliliter.

### Antibodies

Rabbit polyclonal against SARS S1 antibodies (#40150-T62), mouse monoclonal against SARS S1 antibody (#40150-MM02), rabbit polyclonal against SARS-CoV-2 RBD antibodies(#40592-T62), rabbit polyclonal against SARS-CoV-2 S2 antibodies(#40590-T62), rabbit polyclonal against HIV-1 Gag-p24 antibody (11695-RB01) were purchased from Sino Biological Inc. (Beijing, China). Mouse monoclonal anti-FLAG M2 antibody and Mouse monoclonal anti-β-Actin antibody were purchased from Sigma-Aldrich. Integrin β-1 rabbit polyclonal antibody was purchased from Proteintech (Wuhan, China). Alexa flour 488 conjugated rabbit monoclonal His-tag was purchased from Cell Signaling Technology (Danvers, MA, USA). Fluorescein-conjugated goat anti-human IgG (#ZF-0308) was purchased from ZSGB-BIO (Beijing, China). Donkey anti-rabbit IgG (#711-035-152), goat anti-mouse IgG (#115-035-146), rabbit anti-goat IgG (#305-035-003) were purchased from Jackson ImmunoResearch (West Grove, PA, USA).

### Expression and purification of SL-CoV RaTG13, SARS-CoV-2 and SARS-CoV RBDs

Receptor-binding domains (RBDs) of SL-CoV RaTG13, SARS-CoV-2 and SARS-CoV were expressed in Hi5 cells using the Bac-to-Bac baculovirus system (Invitrogen). Briefly, the codon optimized DNA sequences encoding the SL-CoV RaTG13 RBD (residues Arg319-Phe541), SARS-CoV-2 RBD (residues Arg319-Phe541), and SARS-CoV RBD (residues Arg306-Phe527) were inserted into pFastBac (Invitrogen) with an N-terminal gp67 signal peptide and a C-terminal 6 × His tag. The constructs were transformed into DH10Bac competent cells, and the resulting bacmids were transfected into Sf9 cells using Cellfectin II Reagent (Invitrogen) to generate initial virus stock. After amplification, viruses were used to infect Hi5 cells at a density of 2 × 106 cells/ml. The supernatants containing the secreted RBDs were harvested at 60 hrs postinoculation and purified using a Ni-NTA column (GE Healthcare), followed by a Superdex 200 gel filtration column (GE Health care).

### Soluble RBD binding assay

HEK293 cells were transfected with plasmids encoding different ACE2 orthologs (Table S1) by polyetherimide (PEI) (Sigma, St Louis, MO, USA). After 40 hrs incubation, cells were washed with PBS, lifted with PBS containing 1 mM EDTA, and immediately washed twice with PBS with 2% FBS. About 2×10^5^ cells were incubated with 5 μg of soluble RATG13,SARS-CoV-2,or SARS-CoV RBD for 1 hr on ice. After washing three times with PBS with 2% FBS, cells were incubated with rabbit polyclonal anti-6xHis antibody (1:200 dilution) (Shanghai Enzyme-Linked Biotechnology Co., Shanghai, China), followed by incubation with Alexa Fluor 488-conjugated goat anti-rabbit IgG (1:200). Cells were fixed with 1% paraformaldehyde and analyzed by flow cytometry.

### Pseudovirion production and transduction

For pseudotyped virion production, HEK-293 cells were transfected with psPAX2, pLenti-GFP, and plasmids encoding either SARS-CoV-2 S, SARS-CoV S, RaTG13 S, or ZC45 S protein at equal molar ratios by PEI. After 40 hrs of incubation, viral supernatants were harvested and centrifuged at 800 g for 5 min to remove cell debris. For transduction, receptor-expressing cells were seeded into 24-well plates at 30-40% confluence. The next day, cells were inoculated with 500 μl viral supernatant, followed by spin-inoculation at 800g for 30 min. After overnight incubation, cells were fed with fresh media, and cells were lysed with 120 μl of lysis buffer (ratio of medium and Steady-glo (Promega) at 1:1) at 48 hrs postinoculation. The luciferase activities were quantified by using a Modulus II microplate reader (Turner Biosystems, Sunnyvale, CA, USA). All experiments were performed in triplicate and repeated at least twice.

### Detection of S protein by western blot

Briefly, HEK293T cells transfected with plasmids encoding either SARS-CoV, SARS-CoV-2, bat SL-CoV RaTG13, or bat SL-CoV ZC45 S proteins were lysed at 40 hrs post transfection by RIPA buffer (20 mM Tris-HCl pH 7.5, 150 mM NaCl, 1 mM EDTA, 0.1% SDS, 1% NP40, 1× protease inhibitor cocktail). After 30 min of incubation on ice, cell lysate was centrifuged at 12,000g for 10 min at 4°C to remove nuclei. To pellet down pseudovirions, viral supernatants were centrifuged at 25,000 rpm for 2 hrs in a Beckman SW41 rotor at 4°C through a 20% sucrose cushion, and virion pellets were resuspended in 30 μl RIPA buffer. The samples were boiled for 10 min, separated in a 10% SDS-PAGE gel (WB1102, Beijing Biotides Biotechnology, Beijing, China) and transferred to nitrocellulose filter membranes. After blocking with 5% milk, the membranes were blotted with primary antibodies, followed by horseradish peroxidase (HRP) conjugated secondary antibodies (1:5000), and visualized with Chemiluminescent Reagent (Bio-Rad). The primary antibodies used for blotting were polyclonal goat anti-MHV S antibody AO4 (1:2000), polyclonal anti-SARS S1 antibodies T62 (1:2000) (Sinobiological Inc, Beijing, China), mouse monoclonal against SARS S1 antibody MM02 (1:1000) (Sinobiological Inc, Beijing, China), rabbit polycolonal anti-SARS-CoV-2 RBS antibodies (1:1000) (Sinobiological Inc, Beijing, China), rabbit polycolonal anti-SARS-CoV-2 S2 antibodies (1:1000) (Sinobiological Inc, Beijing, China) and anti-FLAG M2 antibody (1:1000) (Sigma, St. Louis, MO, USA), respectively.

### Cell surface protein biotinylation assay

To determine the level of ACE2s of each species on the cell surface, FLAG-tagged ACE2 expressing cells at 80-90% confluence were incubated with PBS containing 2.5 μg/mL EZ-linked Sulfo-NHS-LC-LC-biotin (Thermo-Pierce, #21388) on ice for 30 min after washing with ice-cold PBS. Then, the reaction was quenched by PBS with 100 mM glycine and cells were lysed with RIPA buffer. To pull-down the proteins labeled with biotin, the lysates were incubated with NeutrAvidin beads (Thermo-Pierce, #53150) overnight at 4°C. After washing 3 times with RIPA buffer, samples were resuspended in 30 μl of loading buffer and boiled for 10 min, and the level of ACE2 expression was determined by western-blotting using an anti-FLAG M2 antibody (1:1000). Integrin-β1 were serving as a control.

### Cell-cell fusion assay

HEK293T cells transiently overexpressing the S protein and eGFP were detached by brief trypsin (0.25%) treatment, and overlaid on a 70% confluent monolayer of ACE2 expressing cells at a ratio of approximately one S-expressing cell to three receptor-expressing cells. After 4 hrs of incubation, images of syncytia were captured with a Nikon TE2000 epifluorescence microscope running MetaMorph software (Molecular Devices). All experiments were performed in triplicate and repeated at least three times. Three images for each sample were selected, and the total number of nuclei and the number of nuclei in fused cells for each image were counted. The fusion efficiency was calculated as the number of nuclei in syncytia/total number of nuclei x100.

### Structure modeling

The PDB files of the crystal structures of hACE2/SARS-CoV-2 (6m0j)and hACE2/SARS-CoV (2ajf) and the cryo-EM structure of the RaTG13 spike glycoprotein (6zgf) were downloaded from the RCSB PDB website (www.rcsb.org). Homology models of the receptor binding domain (RBD) of different host ACE2s were built with the Structuropedia web server (mod.farooq.ac). Hot spot residues were predicted with the Hotpoint web server (prism.ccbb.ku.edu.tr/hotpoint)^42,43^. The RBD structures of SARS-CoV-2, SARS-CoV, and RaTG13 were extracted from the pdb files and docked into the homology models with the HADDOCK server (wenmr.science.uu.nl)^44^, using conserved active residues on the interfaces as docking restraints. Docking poses were viewed, aligned, and analyzed with PyMOL software.

## ACKNOWLEDGEMENT

This work was supported by grants from the National Key R&D Program of China (2020YFA0707600 to ZQ), the National Natural Science Foundation of China (31670164 and 31970171 to ZQ), and the CAMS Innovation Fund for Medical Sciences (2016-12M-1-014 to JW)

## Reference

1 Huang, C. et al. Clinical features of patients infected with 2019 novel coronavirus in Wuhan, China. Lancet, doi:10.1016/S0140-6736(20)30183-5 (2020).

2 Ren, L. L. et al. Identification of a novel coronavirus causing severe pneumonia in human: a descriptive study. Chin Med J (Engl), doi:10.1097/CM9.0000000000000722 (2020).

3 Zhou, P. et al. A pneumonia outbreak associated with a new coronavirus of probable bat origin. Nature, doi:10.1038/s41586-020-2012-7 (2020).

4 Zhu, N. et al. A Novel Coronavirus from Patients with Pneumonia in China, 2019. N Engl J Med, doi:10.1056/NEJMoa2001017 (2020).

5 WHO. Coronavirus disease (COVID-2019) situation reports. https://www.who.int/emergencies/diseases/novel-coronavirus-2019/situation-reports/. (2020).

6 Viruses, I. C. o. T. o. Virus Taxonomy: 2019 Release. https://talk.ictvonline.org/taxonomy/. (2019).

7 Masters, P. S. & Perlman, S. in Fields Virology Vol. 1 (eds D.M. Knipe & P.M Howley) Ch. 8, 825–858 (2013).

8 Li, X. et al. Emergence of SARS-CoV-2 through Recombination and Strong Purifying Selection. bioRxiv, doi:10.1101/2020.03.20.000885 (2020).

9 Xiao, K. et al. Isolation of SARS-CoV-2-related coronavirus from Malayan pangolins. Nature 583, 286–289, doi:10.1038/s41586-020-2313-x (2020).

10 Ou, X. et al. Characterization of spike glycoprotein of SARS-CoV-2 on virus entry and its immune cross-reactivity with SARS-CoV. Nature communications 11, 1620, doi:10.1038/s41467-020-15562-9 (2020).

11 Hoffmann, M. et al. SARS-CoV-2 Cell Entry Depends on ACE2 and TMPRSS2 and Is Blocked by a Clinically Proven Protease Inhibitor. Cell 181, 271–280 e278, doi:10.1016/j.cell.2020.02.052 (2020).

12 Lan, J. et al. Structure of the SARS-CoV-2 spike receptor-binding domain bound to the ACE2 receptor. Nature 581, 215–220, doi:10.1038/s41586-020-2180-5 (2020).

13 Shang, J. et al. Structural basis of receptor recognition by SARS-CoV-2. Nature 581, 221–224, doi:10.1038/s41586-020-2179-y (2020).

14 Wang, Q. et al. Structural and Functional Basis of SARS-CoV-2 Entry by Using Human ACE2. Cell 181, 894–904 e899, doi:10.1016/j.cell.2020.03.045 (2020).

15 Yan, R. et al. Structural basis for the recognition of SARS-CoV-2 by full-length human ACE2. Science 367, 1444–1448, doi:10.1126/science.abb2762 (2020).

16 Zhao, X. et al. Broad and differential animal ACE2 receptor usage by SARS-CoV-2. J Virol, doi:10.1128/JVI.00940-20 (2020).

17 Shi, J. et al. Susceptibility of ferrets, cats, dogs, and other domesticated animals to SARS-coronavirus 2. Science 368, 1016–1020, doi:10.1126/science.abb7015 (2020).

18 Oreshkova, N. et al. SARS-CoV-2 infection in farmed minks, the Netherlands, April and May 2020. Euro Surveill 25, doi:10.2807/1560-7917.ES.2020.25.23.2001005 (2020).

19 Wang, L. et al. Complete Genome Sequence of SARS-CoV-2 in a Tiger from a U.S. Zoological Collection. Microbiol Resour Announc 9, doi:10.1128/MRA.00468-20 (2020).

20 Imai, M. et al. Syrian hamsters as a small animal model for SARS-CoV-2 infection and countermeasure development. Proc Natl Acad Sci U S A 117, 16587–16595, doi:10.1073/pnas.2009799117 (2020).

21 Halfmann, P. J. et al. Transmission of SARS-CoV-2 in Domestic Cats. N Engl J Med, doi:10.1056/NEJMc2013400 (2020).

22 Zhai, X. et al. Comparison of Severe Acute Respiratory Syndrome Coronavirus 2 Spike Protein Binding to ACE2 Receptors from Human, Pets, Farm Animals, and Putative Intermediate Hosts. J Virol 94, doi:10.1128/JVI.00831-20 (2020).

23 Li, Y. et al. SARS-CoV-2 and three related coronaviruses utilize multiple ACE2 orthologs and are potently blocked by an improved ACE2-Ig. J Virol, doi:10.1128/JVI.01283-20 (2020).

24 Wrobel, A. G. et al. SARS-CoV-2 and bat RaTG13 spike glycoprotein structures inform on virus evolution and furin-cleavage effects. Nature structural & molecular biology, doi:10.1038/s41594-020-0468-7 (2020).

25 Hu, D. et al. Genomic characterization and infectivity of a novel SARS-like coronavirus in Chinese bats. Emerg Microbes Infect 7, 154, doi:10.1038/s41426-018-0155-5 (2018).

26 Qian, Z., Dominguez, S. R. & Holmes, K. V. Role of the spike glycoprotein of human Middle East respiratory syndrome coronavirus (MERS-CoV) in virus entry and syncytia formation. PLoS One 8, e76469, doi:10.1371/journal.pone.0076469 (2013).

27 Hou, Y. et al. Angiotensin-converting enzyme 2 (ACE2) proteins of different bat species confer variable susceptibility to SARS-CoV entry. Arch Virol 155, 1563–1569, doi:10.1007/s00705-010-0729-6 (2010).

28 Lam, T. T. et al. Identifying SARS-CoV-2-related coronaviruses in Malayan pangolins. Nature 583, 282–285, doi:10.1038/s41586-020-2169-0 (2020).

29 Liu, P. et al. Are pangolins the intermediate host of the 2019 novel coronavirus (SARS-CoV-2)? PLoS Pathog 16, e1008421, doi:10.1371/journal.ppat.1008421 (2020).

30 Zhang, T., Wu, Q. & Zhang, Z. Probable Pangolin Origin of SARS-CoV-2 Associated with the COVID-19 Outbreak. Curr Biol 30, 1346–1351 e1342, doi:10.1016/j.cub.2020.03.022 (2020).

31 Dinnon, K. H. et al. A mouse-adapted SARS-CoV-2 model for the evaluation of COVID-19 medical countermeasures. bioRxiv, doi:10.1101/2020.05.06.081497 (2020).

32 Kim, Y. I. et al. Infection and Rapid Transmission of SARS-CoV-2 in Ferrets. Cell host & microbe 27, 704–709 e702, doi:10.1016/j.chom.2020.03.023 (2020).

33 Richard, M. et al. SARS-CoV-2 is transmitted via contact and via the air between ferrets. Nature communications 11, 3496, doi:10.1038/s41467-020-17367-2 (2020).

34 Yi, C. et al. Key residues of the receptor binding motif in the spike protein of SARS-CoV-2 that interact with ACE2 and neutralizing antibodies. Cell Mol Immunol 17, 621–630, doi:10.1038/s41423-020-0458-z (2020).

35 Lau, S. K. P. et al. Possible Bat Origin of Severe Acute Respiratory Syndrome Coronavirus 2. Emerg Infect Dis 26, 1542–1547, doi:10.3201/eid2607.200092 (2020).

36 Andersen, K. G., Rambaut, A., Lipkin, W. I., Holmes, E. C. & Garry, R. F. The proximal origin of SARS-CoV-2. Nat Med 26, 450–452, doi:10.1038/s41591-020-0820-9 (2020).

37 Boni, M. F. et al. Evolutionary origins of the SARS-CoV-2 sarbecovirus lineage responsible for the COVID-19 pandemic. Nat Microbiol, doi:10.1038/s41564-020-0771-4 (2020).

38 Lee, J. et al. No evidence of coronaviruses or other potentially zoonotic viruses in Sunda pangolins (Manis javanica) entering the wildlife trade via Malaysia. BioRxiv, doi:https://doi.org/10.1101/2020.06.19.158717.

39 Schlottau, K. et al. SARS-CoV-2 in fruit bats, ferrets, pigs, and chickens: an experimental transmission study. Lancet Microbe 1, e218–e225, doi:10.1016/S2666-5247(20)30089-6 (2020).

40 Park, S. J. et al. Antiviral Efficacies of FDA-Approved Drugs against SARS-CoV-2 Infection in Ferrets. mBio 11, doi:10.1128/mBio.01114-20 (2020).

41 Wang, J. et al. Mouse-adapted SARS-CoV-2 replicates efficiently in the upper and lower respiratory tract of BALB/c and C57BL/6J mice. Protein Cell, doi:10.1007/s13238-020-00767-x (2020).

42 Tuncbag, N., Gursoy, A. & Keskin, O. Identification of computational hot spots in protein interfaces: combining solvent accessibility and inter-residue potentials improves the accuracy. Bioinformatics 25, 1513–1520, doi:10.1093/bioinformatics/btp240 (2009).

43 Tuncbag, N., Keskin, O. & Gursoy, A. HotPoint: hot spot prediction server for protein interfaces. Nucleic Acids Res 38, W402–406, doi:10.1093/nar/gkq323 (2010).

44 van Zundert, G. C. P. et al. The HADDOCK2.2 Web Server: User-Friendly Integrative Modeling of Biomolecular Complexes. Journal of molecular biology 428, 720–725, doi:10.1016/j.jmb.2015.09.014 (2016).

